# Population dynamics of *Escherichia coli* in the gastrointestinal tracts of Tanzanian children

**DOI:** 10.1101/294934

**Authors:** Taylor K. S. Richter, Tracy H. Hazen, Diana Lam, Christian L. Coles, Jessica C. Seidman, Yaqi You, Ellen K. Silbergeld, Claire M. Fraser, David A. Rasko

## Abstract

The stability of the *Escherichia coli* populations in the human gastrointestinal tract are not fully appreciated, and represent a significant knowledge gap regarding gastrointestinal community structure, as well as resistance to incoming pathogenic bacterial species and antibiotic treatment. The current study examines the genomic content of 240 *Escherichia coli* isolates from children 2 to 35 months old in Tanzania. The *E. coli* strains were isolated from three time points spanning a six month time period, with or without antibiotic treatment. The resulting isolates were sequenced, and the genomes compared. The findings in this study highlight the transient nature of *E. coli* strains in the gastrointestinal tract of children, as during a six-month interval, no one individual contained phylogenomically related isolates at all three time points. While the majority of the isolates at any one time point were phylogenomically similar, most individuals did not contain phylogenomically similar isolates at more than two time points. Examination of global genome content, canonical *E. coli* virulence factors, multilocus sequence type, serotype, and antimicrobial resistance genes identified diversity even among phylogenomically similar strains. There was no apparent increase in the antimicrobial resistance gene content after antibiotic treatment. The examination of the *E. coli* from longitudinal samples from multiple children in Tanzania provides insight into the genomic diversity and population variability of resident *E. coli* within the rapidly changing environment of the gastrointestinal tract.

**Importance:** This study increases the number of resident *Escherichia coli* genome sequences, and explores *E. coli* diversity through longitudinal sampling. We investigate the genomes of *E. coli* isolated from human gastrointestinal tracts as part of an antibiotic treatment program among rural Tanzanian children. Phylogenomics demonstrates that resident *E. coli* are diverse, even within a single host. Though the *E. coli* isolates of the gastrointestinal community tend to be phylogenomically similar at a given time, they differed across the interrogated time points, demonstrating the variability of the members of the *E. coli* community. Exposure to antibiotic treatment did not have an apparent impact on the *E. coli* community or the presence of resistance and virulence genes within *E. coli* genomes. The findings of this study highlight the variable nature of bacterial members of the human gastrointestinal tract.

## Introduction

*Escherichia coli* in the human gastrointestinal tract is often recognized as an important source of disease (1, 2). As the causative agent of over 2 million deaths annually due to diarrhea (3, 4), as well as millions of extraintestinal infections (5), its categorization as a pathogen is not unwarranted. Particularly in developing countries, the consequences of diarrheal *E. coli* is substantial among children under five years old, who incur the majority of infections and deaths (3) and whose rapidly developing microbiomes can be impacted by frequent bouts of disease and treatment (6, 7). Yet, *E. coli* are the dominant aerobic organism in the human gastrointestinal tract, identified in greater than 90% of humans, and many other large mammals, often reaching concentrations up to 10^9^ CFU per gram of feces (8) without causing disease. In this role as a resident organism in healthy hosts, they are thought to have critical roles in digestion, nutrition, metabolism, and protection against incoming enteric pathogens (9-12). Despite the importance and involvement of *E. coli* in human health, studies of their role as native, non-pathogenic members of the human gastrointestinal microbiome are poorly represented among genome sequencing, comparative analysis efforts, and functional characterization.

Most studies of bacterial genomics have focused on pathogenic isolates over a limited time frame. *E. coli* genomic studies are no exception, having concentrated on sequencing single isolates, from single time points, and on samples related to a clinical presentation, such as diarrhea or urinary tract infection (10, 13-17). There have been fewer than five genomes sequenced of non-pathogenic *E. coli*, in addition to a limited number of isolates from the feces of individuals that do not have diarrhea (10, 17-20). To date the genomic examination of longitudinal isolates is lacking, thus hindering the ability to explore the diversity of *E. coli* isolates both within-host and across time. Most studies of resident *E. coli* were completed prior to ready access to sequencing technologies (11). An exception is Stoesser *et al.* (18), which identified multiple isolates in single-host samples using single nucleotide polymorphism (SNP) level analyses, leaving much to be learned about *E. coli* genomic diversity within and between human hosts over longitudinal sampling.

A population-based longitudinal cohort study, PRET+ (Partnership for the Rapid Elimination of Trachoma, January to July 2009), provided a unique opportunity to examine both the diversity and dynamics of the *E. coli* isolates in the human gastrointestinal tract among children in Tanzania (21, 22). In the PRET+ study, Seidman *et al*. investigated the effects of mass distribution of azithromycin on antibiotic resistance of resident *E. coli* (21, 22). *E. coli* were isolated from fecal swabs obtained from children 2 to 35 months old living in rural Tanzania, half of whom were given a single oral prophylactic azithromycin treatment for trachoma (an infection of the eye caused by *Chlamydia trachomatis*). *E. coli* isolates from this cohort were selected for genome sequencing and comparative analyses to investigate the within-subject and longitudinal diversity of *E. coli* isolates in children (Table S1). Up to three isolates per individual, from each of three time points spanning six months, were collected in the PRET+ study, providing up to nine potential isolates from each subject for examination (Figure 1).

**Figure 1:**
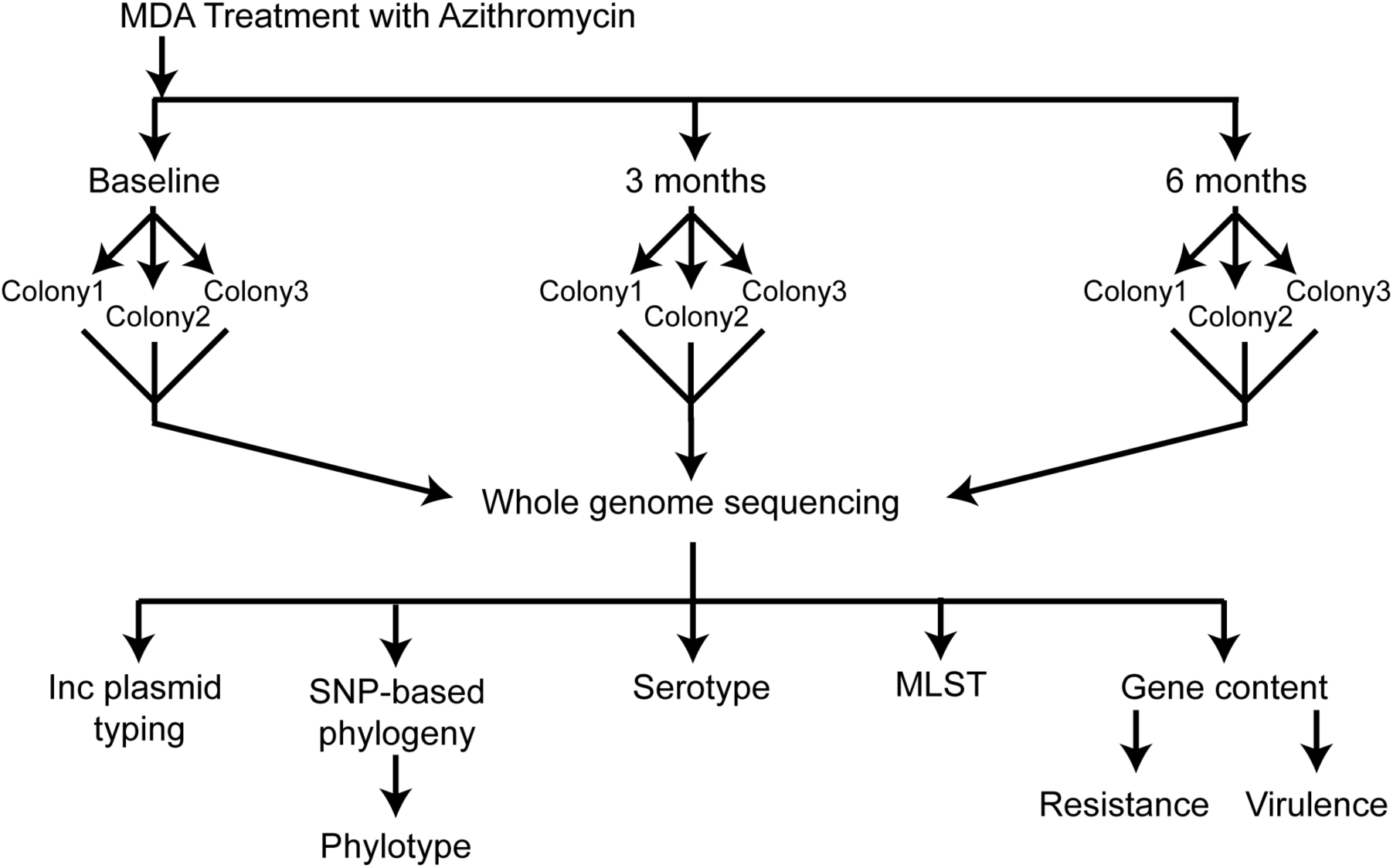
Overall study design. The overall design of the study highlighting the sampling of up to three distinct colonies on three time points, one of which, termed the baseline occurs prior to the administration of antibiotics in half of the subjects.

Samples from the current study provide insight into *E. coli* diversity within a subject over several time points. While other studies have examined resident *E. coli* in children in developing countries, they limited their focus to using PCR and *in vitro* lab techniques to identify a limited set of canonical virulence genes and determine resistance profiles of the isolated strains (23-25). In addition to the virulence- and resistance-associated gene content, the current study demonstrates previously uncharacterized diversity among *E. coli* isolates from the human gastrointestinal tract on a whole genome level within and across sampling periods. This work represents the most comprehensive longitudinal genomic study of resident *E. coli* within the human gastrointestinal tract and expands knowledge of the non-pathogen gut flora by increasing the available genome sequences of resident *E. coli* and highlighting the dynamic nature of the *E. coli* community.

## Results

### Selection of E. coli strains for genome sequencing

A total of 247 *E. coli* isolates from 30 subjects (17 male and 13 female as shown in Figure 2) in the study by Seidman et al. (21, 22) were selected for DNA extraction and genome assembly, based on the criteria that these subjects contributed the most complete longitudinal collection of isolates (i.e. the greatest number of subjects with the greatest number of possible isolates). Of these, 240 isolates provided acceptable sequence quality to generate genome assemblies with a genome size and GC-content that is characteristic of *E. coli* to be analyzed using comparative genomics. The average genome size was 5.17 Mb (range 4.46 to 5.81 Mb) with a 50.69% GC (range 50.21 to 51.04%), similar to other known *E. coli* genomes (Table S1). Of the 240 isolates, 120 isolates were from the subjects that received the antibiotic treatment of single oral dose of prophylactic azithromycin, and 120 isolates from subjects in the non-treatment (control) group (Table S1 and Fig.2).

**Figure 2:**
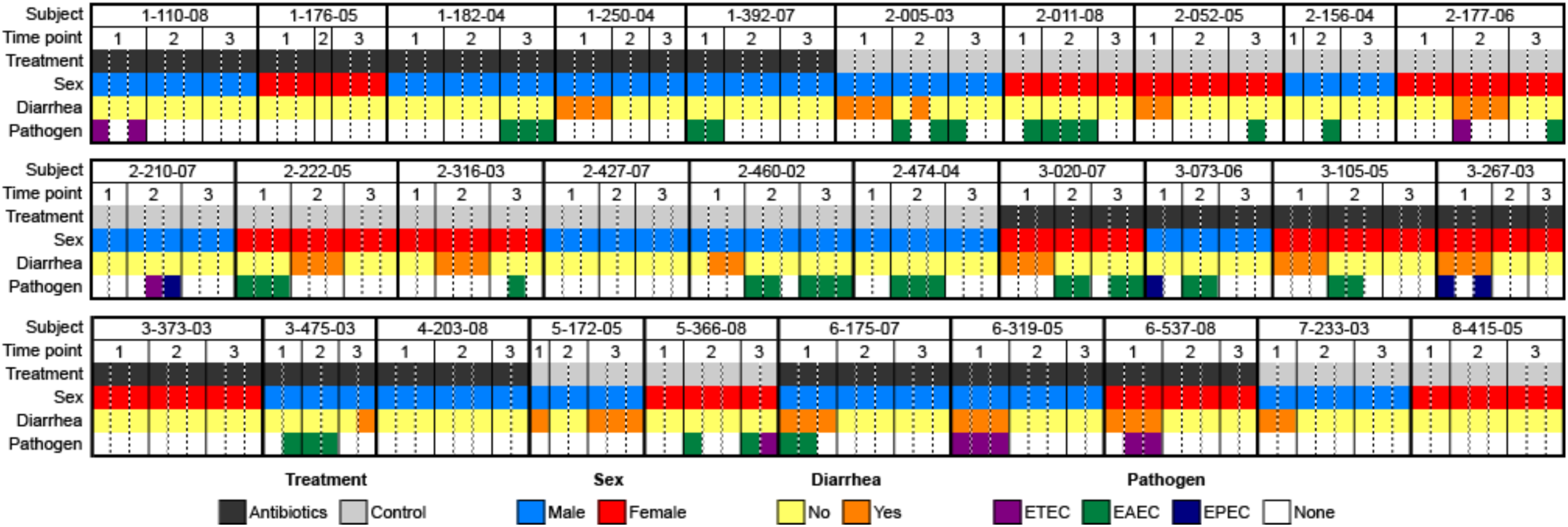
Isolate metadata. Summary of metadata showing time point of isolation, treatment group, host sex, clinical presentation, and the identification of pathogenic markers for ETEC, EAEC, or EPEC pathotypes for each isolate by subject. Further details in Table S1

### Subject clinical state and E. coli pathotype identification

There were 17 instances in which subjects had active diarrhea at the time of sample collection (12 instances occurred at the baseline time point), yielding 46 isolates from diarrheal conditions (21, 22), 23 each from the antibiotic treatment and control groups. All cases of diarrhea were identified in children under the age of 2 at baseline. Only 16 of these isolates (34.8%) contained canonical virulence factors belonging to the EPEC, ETEC, or EAEC pathotypes (Fig. 2), as determined by sequence homology searches of canonical virulence genes in the assembled genomes. In most cases, observed diarrhea could not be associated with a prototypically virulent *E. coli* in this data set. Other sources of diarrhea were not investigated.

An additional 61 isolates from 19 individuals contained canonical *E. coli* virulence factors, but were not obtained from samples taken during an active diarrheal event. These data indicate that the presence of a potentially virulent *E. coli* does necessarily result in clinical presentation of diarrhea. Overall, in our dataset there was no evidence of an association between diarrheal cases and incidence of isolates containing canonical *E. coli* virulence factors.

### Phylogenomic analysis

Phylogenomic analysis of the isolates identified a diverse population of *E. coli* within the gastrointestinal community of these children. A phylogenetic tree of the 240 isolates from this study plus 33 reference *E. coli* and *Shigella* genomes (Table S2) was used to assess the genomic similarity of the isolates from a single subject both within and across time points, as well as between subjects over the study period (Fig. 3). The SNP-based phylogenomic analysis of the draft and reference genomes identified 304,497 polymorphic single nucleotide genomic sites. The isolates from the current study were identified in the established *E. coli* phylogroups: A (132 isolates), B1 (62 isolates), B2 (24 isolates), D (17 isolates), and E (2 isolates) (Fig. 3, Table S1). Additionally, three isolate genomes (isolates 1_176_05_S3_C2, 2_011_08_S1_C1, and 2_156_04_S3_C2) fell into cryptic clades located outside of the established *E. coli* phylogroups. The distributions of the *E. coli* isolates in each of these phylogroups were not associated with any of the clinical parameters associated with these isolates.

**Figure 3.**
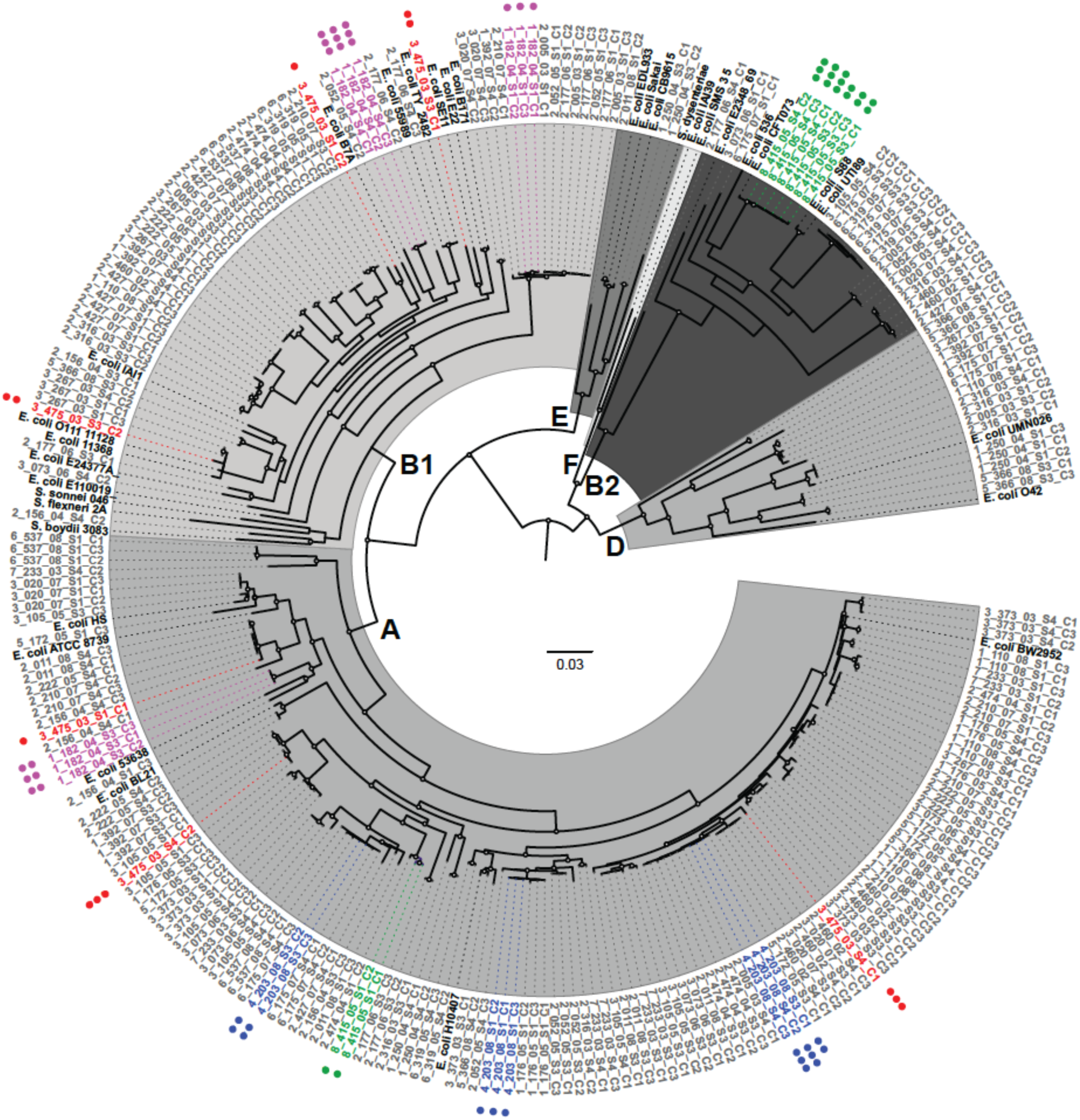
Phylogenomic analysis of *E. coli* isolates in study. **A)** A whole-genome phylogeny of the isolate sequences and reference *E. coli* and *Shigella* genomes (shown in black) highlighting examples of diversity among subject-specific isolates within and across time points. The scale bar indicates the approximate distance of 0.03 nucleotide substitutions per site Nodes with bootstrap values of greater than 90 are marked with a circle. Examples of isolates from subjects that demonstrate the greatest (3_475_03) and least (4_203_08, 8_415_05, 1_182_04) amount of diversity are highlighted: 3_475_03 in red, 4_203_08 in blue, 8_415_05 in green, and 1_182_04 in purple. The number of dots denote the sample number from which the isolate was obtained. *E. coli* phylogroups are labeled. Full figure with all subjects is presented in Figure S1.

To further investigate the *E. coli* diversity of an individual subject at a given time, we analyzed the phylogenetic groupings of isolates from each subject at each time point. Most isolates from an individual at a single time point group together within a single phylogenomic lineage, where a lineage is defined as a terminal grouping of isolates (54.4%; 49 of the 90 same-subject time points). One third (35.5%; 32/90 of the same-subject time point isolates) fell into two distinct lineages, and in 10% (9/90 time points), all isolates belonged to a distinct lineage (Table 1). Overall, these data suggest that while there is considerable diversity among the isolates from many subjects, in over half of them, the population of *E. coli* at a given time point displays limited phylogenomic variation.

**Table 1.** Summary of isolate diversity within subject and within time point across several diversity measurements

| Subject ID | Treatment | Isolate Phylogenomics |  | Resistance |  |  | Virulence |  |  | Phylogroup |  | MLST |  | Serotype |  |
| --- | --- | --- | --- | --- | --- | --- | --- | --- | --- | --- | --- | --- | --- | --- | --- |
|  |  | No. isolates from subject | No. of clades in subject* | No. resistance clades* | Isolates single resistance superclade | Similar distribution to phylogeny* | No. virulence gene clades* | Similar distribution to phylogeny* | Similar distribution to resistance genes* | No. phylogroups in subject* | Similar distribution to phylogeny* | No. sequence types in subject* | Similar distribution to phylogeny* | No. serotypes in subject* | Similar distribution to phylogeny* |
| 1_110_08 | MDA | 9 | 5 | 5 | No | No | 5 | No | Yes | 3 | No | 3 | No | 5 | Yes |
| 1_176_05 | MDA | 8 | 4 | 4 | No | Yes | 4 | Yes | Yes | 2 | No | 3 | No | 4 | Yes |
| 1_182_04 | MDA | 9 | 3 | 5 | No | No | 3 | Yes | No | 2 | No | 3 | Yes | 3 | Yes |
| 1_250_04 | MDA | 7 | 3 | 2 | Yes | No | 3 | Yes | No | 3 | Yes | 3 | Yes | 3 | Yes |
| 1_392_07 | MDA | 8 | 4 | 5 | No | No | 4 | Yes | No | 3 | No | 4 | Yes | 4 | Yes |
| 3_020_07 | MDA | 8 | 4 | 4 | No | Yes | 4 | Yes | Yes | 5 | Yes | 5 | Yes | 4 | No |
| 3_073_06 | MDA | 7 | 5 | 5 | No | Yes | 4 | No | No | 3 | No | 6 | No | 7 | No |
| 3_105_05 | MDA | 9 | 7 | 6 | No | Yes | 7 | Yes | No | 3 | No | 5 | Yes | 5 | Yes |
| 3_267_03 | MDA | 7 | 6 | 6 | No | Yes | 6 | Yes | Yes | 2 | No | 6 | No | 7 | Yes |
| 3_373_03 | MDA | 9 | 4 | 4 | No | Yes | 3 | No | No | 3 | No | 6 | Yes | 6 | Yes |
| 3_475_03 | MDA | 6 | 6 | 5 | No | No | 6 | Yes | No | 2 | No | 6 | Yes | 6 | Yes |
| 4_203_08 | MDA | 8 | 3 | 5 | No | No | 3 | No | No | 2 | No | 4 | Yes | 4 | Yes |
| 6_175_07 | MDA | 9 | 4 | 5 | No | Yes | 5 | Yes | Yes | 3 | No | 6 | Yes | 5 | No |
| 6_319_05 | MDA | 8 | 3 | 5 | No | No | 3 | Yes | No | 3 | No | 5 | No | 7 | No |
| 6_537_08 | MDA | 8 | 3 | 5 | No | No | 3 | Yes | No | 3 | No | 4 | Yes | 5 | No |
| 2_005_03 | No-MDA | 9 | 5 | 7 | No | No | 6 | Yes | No | 2 | No | 4 | Yes | 4 | Yes |
| 2_011_08 | No-MDA | 8 | 6 | 5 | No | No | 6 | Yes | No | 3 | No | 4 | Yes | 4 | Yes |
| 2_052_05 | No-MDA | 8 | 5 | 4 | No | Yes | 5 | No | No | 3 | No | 5 | No | 5 | No |
| 2_156_04 | No-MDA | 7 | 7 | 5 | No | Yes | 6 | No | No | 2 | No | 7 | Yes | 7 | Yes |
| 2_177_06 | No-MDA | 9 | 6 | 5 | No | No | 7 | No | No | 3 | No | 6 | Yes | 6 | Yes |
| 2_210_07 | No-MDA | 8 | 6 | 6 | No | Yes | 5 | No | No | 1 | No | 3 | No | 4 | Yes |
| 2_222_05 | No-MDA | 9 | 4 | 5 | No | No | 4 | Yes | No | 2 | No | 6 | Yes | 6 | Yes |
| 2_316_03 | No-MDA | 8 | 6 | 7 | No | No | 5 | No | No | 1 | No | 2 | No | 3 | Yes |
| 2_427_07 | No-MDA | 8 | 5 | 4 | No | Yes | 6 | No | No | 1 | No | 4 | Yes | 4 | Yes |
| 2_460_02 | No-MDA | 9 | 4 | 4 | No | Yes | 4 | Yes | Yes | 3 | No | 6 | No | 5 | Yes |
| 2_474_04 | No-MDA | 8 | 4 | 3 | No | No | 3 | No | Yes | 3 | No | 4 | Yes | 4 | Yes |
| 5_172_05 | No-MDA | 6 | 4 | 3 | No | No | 4 | Yes | No | 3 | Yes | 4 | No | 3 | Yes |
| 5_366_08 | No-MDA | 7 | 5 | 5 | No | Yes | 4 | No | No | 2 | No | 3 | Yes | 3 | Yes |
| 7_233_03 | No-MDA | 8 | 5 | 5 | Yes | Yes | 5 | Yes | Yes | 1 | No | 4 | No | 6 | No |
| 8_415_05 | No-MDA | 8 | 2 | 3 | No | No | 2 | Yes | No | 2 | Yes | 3 | No | 2 | Yes |

\* Further details are provided in Supplemental Table 3

These *E. coli* populations were variable over time, showing increased *E. coli* diversity in each subject when observed over the multiple time points. Same-subject isolates from different time points reside in distinct phylogenomic lineages in 93.3% (28/30) of subjects. Only two subjects had isolates from multiple time points that occupied the same clade. Subject 4_203_08 had one 3-month isolate that was most similar to the two 6-month isolates (Fig. 3). Additionally, subject 8_415_05 had all of the 3-month and 6-month isolates belonging to the same phylogenomic lineage (Fig. 3). Similarly, for subject 2_052_05, all 3-month isolates and one 6-month isolate are in neighboring lineages, suggesting a close phylogenetic relationship.

Only three subjects (1_182_04, 1_250_04, 6_319_05) had a single phylogenomic clade of isolates at each of the three time points (illustrated in Fig. 3 and detailed in Table S3), suggesting colonization by a single dominant clone at any one time point, but dynamic *E. coli* populations between each of the time points. In contrast, all isolates from subject 3_475_03 were phylogenomically distinct (Fig. 3). Additionally, the isolates from eight subjects (26.7%) are represented in at least six distinct phylogenetic groups (Table 1). To our knowledge, this level of phylogenomic diversity of *E. coli* in the human gastrointestinal tract over relatively short time periods has not been previously reported.

### Multilocus sequence typing and molecular serotyping

The genomes in this study comprise a combined total of 87 sequence types (STs) (Table S1). The most common ST was ST10, which was represented by 40 of the *E. coli* genomes, while 40 additional STs occurred only once (Table S1). Only five isolates were from ST131, which has been demonstrated to be associated with the spread of antimicrobial resistance (PMID 24694052). There was, on average, 1.5 (range 1-3) STs among isolates from a subject at a single time point, and an average of 4.4 (range 2-7) STs per subject across all time points. Since the total number of available isolates per subject varied, the values were normalized per the number of isolates, revealing an average of 2 (range 1-4) isolates per sequence type and mimicking the diversity observed in the phylogenetic analyses (Fig. 4, Table S3).

**Figure 4:**
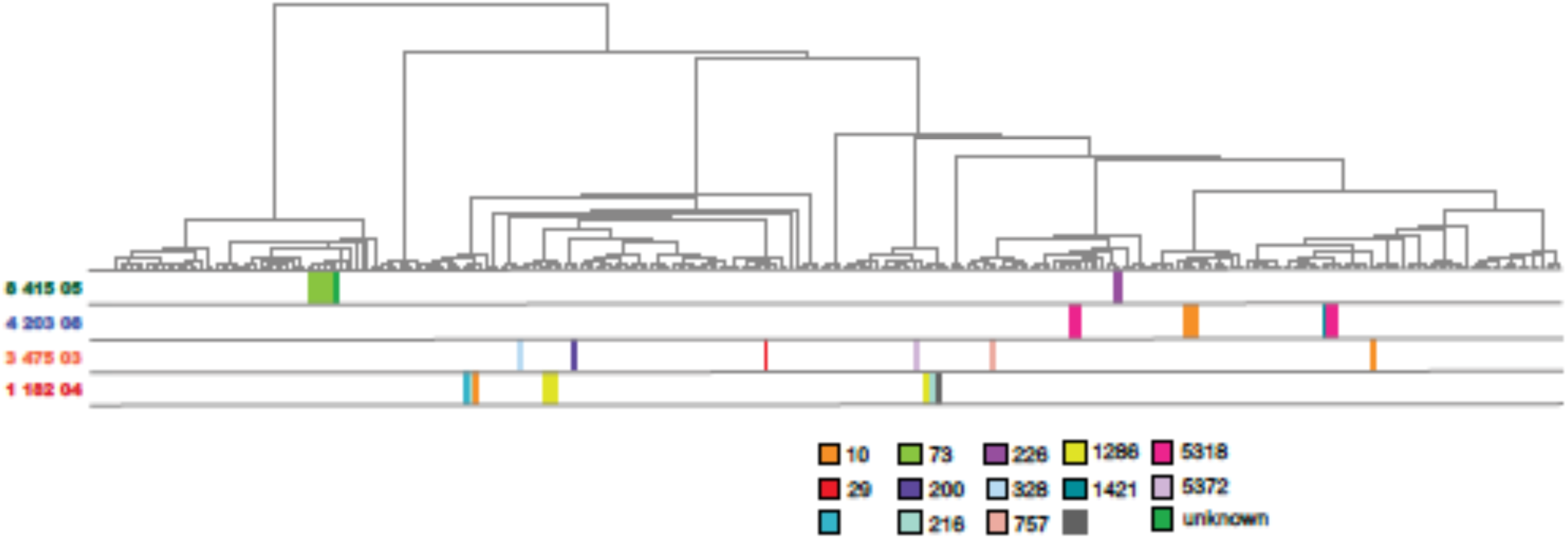
Phylogenomic distribution of sequence types of isolates from select subjects. A cladogram of the phylogeny highlighting relative positions of genomes of isolates from selected subjects with MLST sequence types shown in colored blocks corresponding to the sequence type as shown in the legend. Selected example subjects highlight low diversity within time points but high diversity across time (subject 1_182_04), high diversity within and across time (3_475_03), intermediate diversity across time (4_203_08), and low diversity across time (8_415_05).

Similar to MLST, serotype analyses (26) reflect the diversity observed in the phylogenomic analysis (Table S3). The 240 isolates represent a combined total of 106 O:H serotypes, with 54 of them only occurring once in the dataset, making serotype a finer-scale measure of diversity than MLST. There are an average of 1.63 (range 1-3) different serotypes in isolates from the same time point and 4.7 (range 2-7) serotypes in a subject across all time points. The O, H, or either serotypes could not be predicted in 33 isolates (Table S1). *In silico* analyses were unable to distinguish between some serotypes in an additional 58 isolates (Table S1). This left 149 isolates that could be unambiguously assigned a single serotype (Table S1).

Nearly all isolates that shared a serotype also shared an MLST sequence type and phylogroup (Table S1). There are five examples (excluding those isolates in which the serotype could not be unambiguously differentiated) where MLST, serotype, and phylogroup were not congruent (Table S4), suggesting molecular variation and strain differentiation could not be detected by a single method alone. The combination of these detailed molecular methods could add nuance to diversity measurements in closely related strains.

### Genome content using LS-BSR

Variations in genome content further demonstrated the diversity of the *E. coli* isolate genomes both within and between time points. Using the LS-BSR analysis (27) and an ergatis-based annotation pipeline, a gene content profile was determined which identified 32,950 genes in the pangenome of the 240 isolate genomes. More than 3,000 genes in any single genome was comprised of genes that vary between genomes, leaving only approximately 2000 genes in the conserved core, as has been previously identified (10, 17). This level of variation is true even among the isolates from subject 8_415_05 in which the isolates from the 3-month and 6-month time points group together phylogenetically, and are of the same MLST sequence type. In this case, each isolate contains an average of 220 (range 95-259) variable genes. Given the level of diversity suggested by the variability of the gene content, more detailed SNP analyses, as previously preformed by Stoesser (18) were deemed unnecessary.

### Antibiotic resistance associated gene profiles

The antibiotic treatment of half of the children in this study provided a unique opportunity to investigate the impact of antibiotic treatment on the prevalence and maintenance of antibiotic resistance genes in the *E. coli* community at 3 and 6 months after administration. Antibiotic resistance genes were investigated in the isolate genomes using 1,371 genes from the Comprehensive Antibiotic Resistance Database (CARD) (28). The resistance gene profiles (assortment of present/absent genes) for each isolate were used to create a cladogram to investigate the relationships among isolates by time and by subject (Fig. S2). These relationships were then compared to those in the phylogenetic groupings as well as in the cladogram of virulence gene profiles (Table S5, Figure S3). Similar clustering patterns were identified between the whole genome phylogeny or virulence gene presence and resistance gene-based analysis 74% of the time at each time point, and 37% (phylogeny) or 27% (virulence) of the time for each subject as a whole (Table 1).

There was no significant change in number or type of resistance-associated genes over time, regardless of antibiotic treatment or isolation time point. As subjects were treated with azithromycin, a macrolide, genes conferring resistance to macrolides were investigated in greater detail (Table S6). Macrolide resistance genes were identified in only 19% (46 of the 240) isolates (Table 2) and based on a logistic regression model, there is no evidence to suggest that either time point or antibiotic treatment were significantly associated with macrolide resistance genes (p >0.05 for antibiotic treatment adjusted for time point, for time point adjusted for antibiotic treatment, and overall antibiotic treatment). Isolates from nearly half of the subjects had no known macrolide resistance genes (46.67% antibiotic treatment, 40% control). Based on these results, exposure to a single large dose of azithromycin did not lead to a significant change in the number of known antimicrobial resistance genes or macrolide resistance genes among these *E. coli* populations.

**Table 2:**
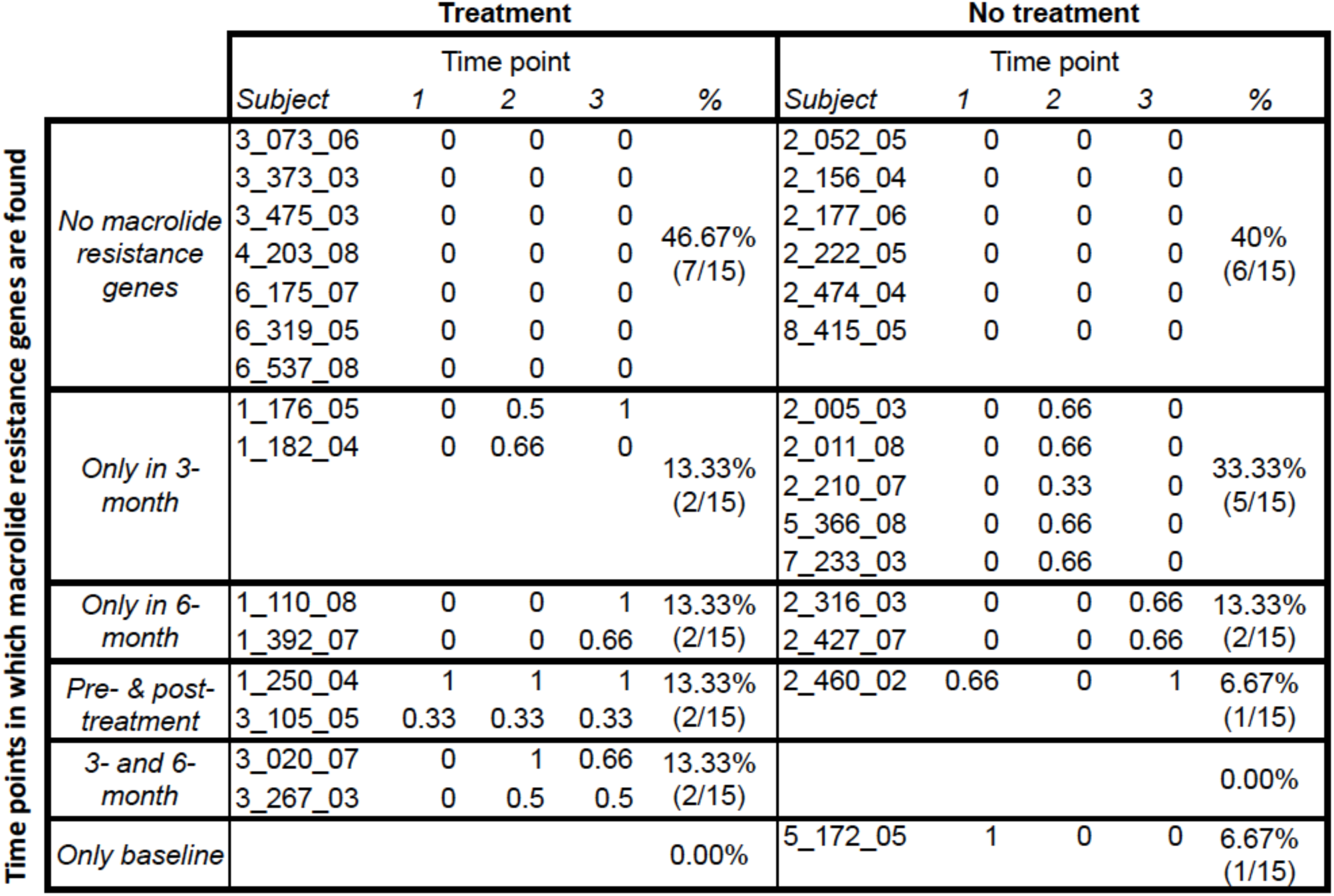
Summary of macrolide resistance gene presence by treatment group and time point. The proportion of isolates in which a macrolide resistance gene was identified is shown for each time point. Subjects are separated in to treatment groups and categorized based on the time points in which macrolide resistance genes are identified. Percentages reflect the proportion of subjects that fall in to each macrolide resistance gene category within treatment groups.

## Discussion

This study represents a detailed examination of the genomic diversity of *Escherichia coli* isolates obtained from longitudinal samples obtained from the gastrointestinal tract of children. An overall trend identified in this study is that the identified *E. coli* from the human gastrointestinal tract are diverse not just between subjects, but within the same subject over time. The *E. coli* genomes sequenced in this study, were selected based on the greatest number of longitudinal isolates per subject and include members of all five of the traditional *E. coli* phlyogroups, as well as 87 different MLST sequence types, and 106 serotypes. The isolates in this study were most frequently of the A or B1 phylogroups, unlike a previous study by Gordon et al (29) in which greater than 70% of the isolates obtained were either from phylogroup B2 or D. This observed difference may be due to differences in sample acquisition (stool swab versus biopsy), or differences in the study participants. The Gordon et al (29) study, obtained samples from adults, the majority (72.5%, 50/69) of whom were diagnosed with either Crohn’s disease or ulcerative colitis, which would likely impact the immune status of the gastrointestinal tract, and potentially alter the bacterial community structure. In contrast our study participants were children under the age of 5, and, other than a few that displayed diarrhea of an unknown source, were considered to be relatively healthy. This study, by using a combination of molecular methods, including whole genome sequencing, enhances the understanding that the *E. coli* in the human gastrointestinal tract is variable and diverse.

Approximately half of *E. coli* isolates in an individual appear phylogenomically and phenotypically similar at any given time point; however, even isolates that appeared clonal based on MLST, phylogroup or serotype still contain unique genomic regions. Gene content analyses revealed variation between isolates thought to be clonal by each of the other methods. However, between time points, the prevalent *E. coli* clones from individual subjects were variable. Only two subjects (4_203_08 and 8_415_05) had isolates that were closely related based on the phylogenomic and other molecular data, at more than one time point (Fig. 3, Fig.4). The more common observations were that distinct and prevalent isolates were present at each 3-month sampling interval.

Previous studies of the variability of *E. coli*, using non-genome sequencing methods, have also identified multiple isolates within a single host, reporting up to 4 *E. coli* genotypes in adult human gastrointestinal studies (18, 29). The findings in this study are similar in that it has identified a number of *E. coli* isolates that are genomically and molecularly different in the subjects at each time, and between time points. While it is possible, and likely, that in the current study less prevalent *E. coli* isolates were not captured at some of the sampling time points, we assume that there was little bias in the selection of the isolates, and that the relative isolate abundance in culture reflects the relative abundance in the feces at the time of sampling. The current study likely still underestimates the *E. coli* diversity in the examined subjects.

Dynamic populations within the human gastrointestinal tract have been previously suggested as an explanation for observations of variable clones in *E. coli* diversity studies (30), but the necessary longitudinal genomic studies were lacking. This study begins to address that deficiency. The observed within-patient and longitudinal diversity of *E. coli* isolates could be a function of age, as all of the subjects in this study were less than three years of age, and thus the diversity could be a result of natural introduction of new exposure to foods, as well as immune system and microbiome development (31, 32). It has been demonstrated that intra-host *E. coli* diversity is greatest in tropical regions where hygiene may play a role and that *E. coli* density in the gastrointestinal tract is altered most significantly in the first two years of a child’s life (11, 33), therefore, it is unclear how well these results correlate with *E. coli* diversity in adults or in other geographic regions. It is thought that the infant microbiome is not established until about three years of age (34), however the detailed longitudinal infant microbiome studies are currently lacking. Future longitudinal studies that include sampling subjects from multiple age groups will be necessary to fully appreciate levels of bacterial population diversity and dynamics present across host populations of all age groups.

Virulence and resistance-associated gene analyses in this study confirm that genomic analyses of single isolates are imperfect predictors of clinical phenotypes, as several isolates harbored canonical *E. coli* virulence genes, classically identifying them as enteric pathogens, but were present in subjects not displaying clinical symptoms, The converse is also possible, in that *E. coli* strains may not contain traditional virulence factors, but be obtained from a diarrheal sample, as has been highlighted in the recent GEMS studies (35, Platts-Mills 2015). There are many potential explanations for these observations which include: 1) the subjects have been previously exposed to these bacteria, and thus, have an established immunity, 2) the organisms are not pathogenic in the context of other host factors, including the host microbiota, 3) additional necessary virulence factors are absent in these isolates, or 4) the virulence factors are present but not expressed by the bacterium. Unfortunately, detailed immunological, microbiota or transcriptional data are not available on the current samples, so the impacts of these factors on pathogenicity cannot be determined conclusively. Whole genome analyses have led to increasing recognition that virulence genes and phylogeny are associated attributes in microbial pathogen genome and suggests that there may be an optimal combination of chromosomal and virulence associated features that results in maximal virulence, survival or transmission (36-39). This may also be true of the success of a commensal isolate in the community (40).

This study adds significantly to the number of available *E. coli* genomes that were not selected for based on pathogenic traits, a group that has been traditionally underrepresented in the sequencing of this species. The scientific community is still in the early stages of understanding gastrointestinal tract microbial ecology and the role that the resident bacteria, including *E. coli*, play in microbiome stability and function. The current study demonstrates that at the genomic level, the community of *E. coli* in the human infant gastrointestinal tract is diverse and variable over time. Further studies on human populations from different geographic areas, as well as other age groups, are required to determine if *E. coli* communities would stabilize as a person approaches adulthood, or whether the community diversity of *E. coli* regularly changes depending on the development of the immune system, as well as many other exposures within the gastrointestinal tract.

## Materials and Methods

### Isolate selection

*E. coli* isolates in this study were selected from isolates collected in Seidman *et al.* (21). The PRET+ study was a 6-month, study designed to assess the ancillary effects on pneumonia, diarrhea and malaria in children following mass distribution of azithromycin for trachoma control. The study was conducted in 8 communities in the `Kongwa, a district located in rural central Tanzania on a semiarid highland plateau with poor access to drinking water. The district has a total population of approximately 248,656, comprising mostly herders and subsistence farmers. The Tanzanian government stipulates that villages with trachoma prevalence ≥10% receive annual mass distribution of azithromycin. On survey, 4 villages found eligible for antibiotic treatment became the PRET+ treatment villages and 4 neighboring ineligible communities were included as controls. The study methods and results detailing the impact of antibiotic treatment on pneumonia and diarrhea morbidity, and antibiotic-resistant *Streptococcus pneumoniae* carriage were published previously (41-43).

The selected *E. coli* isolates were chosen to represent individuals with the most complete longitudinal sample sets from the PRET+ *E. coli* sub-study. Isolates were obtained from 30 individuals between 2-35 months of age, living in 8 villages in the same rural area of Tanzania. Half of these individuals received antibiotic treatment, while the other half (control) received no antibiotic treatment. These isolates were cultured from fecal samples collected at three time points (Figure 1 and Table S1): a baseline prior to antibiotic treatment, three months post-treatment, and six months post-treatment, with corresponding time points in the untreated controls. At each time point, up to three *E. coli* colonies per individual were selected for sequencing and subsequent comparative analyses.

### Bacterial growth and isolation

*E. coli* colonies were obtained as described in Seidman *et al* (21, 22). Briefly, fecal swabs were streaked on MacConkey agar (Difco) and grown overnight at 37°C. Three lactose fermentation (LF) positive colonies were inoculated on nutrient agar stabs and grown overnight at 37°C. *E. coli* isolates were identified as those colonies which were LF-positive, indole-positive (DMACA Indole Reagent droppers, BD), and citrate-negative (Simmons citrate agar slants). Isolates were transferred to Luria broth for overnight growth at 37°C with shaking. *E. coli* cultures were frozen with 10% glycerol and stored at -80°C.

### Genome sequencing and assembly

Genomic DNA was extracted using standard methods (16) and sequenced on the Illumina HiSeq 2000 platform at the Genome Resource Center at the University of Maryland School of Medicine, Institute for Genome Sciences. The resulting 100bp reads were assembled as previously described (36, 38). The assembly details and corresponding GenBank accession numbers are provided in Table S1.

### Identification of predicted pathogen isolates

Isolate genomes were interrogated for the presence of pathotype-specific virulence factor genes using LS-BSR and are derived from a similar *E. coli* typing schema used in the MAL-ED studies (44). The nucleotide sequence for each factor or resistance gene was aligned against all sequenced genomes with BLASTN (45) in conjunction with LS-BSR (27). Genes with a BSR value ≥0.80 were considered highly conserved and present in the isolate examined. The targeted virulence factors are as follows: ETEC heat stable enterotoxin (estA147) or ETEC heat labile enterotoxin (eltb508) identifying the isolate as being enterotoxigenic *E. coli* (ETEC); the a*ggR*-activated island C (aic215) or EAEC ABC transporter A (aata650) genes which are common diagnostic markers for enteroaggregative *E. coli* (EAEC) (46, 47); and the major subunit of the bundle-forming *pilus (bfpA*) (bfpa300) or intimin genes (eae881) which are indicative of enteropathogenic *E. coli* (EPEC) (36).

### Phylogenomic analysis

A total of 273 genomes were used in the phylogenomic analyses: the 240 assembled in this study, in addition to a collection of 33 *E. coli* and *Shigella* reference genomes from GenBank (Table S2). Single nucleotide polymorphisms (SNPs) in all genomes were detected relative to the completed genome sequence of commensal isolate *E. coli* HS (phylogroup A) using the *In Silico* Genotyper (ISG) v.0.12.2 (48), which uses MUMmer v.3.22 (49) for SNP detection. Analysis with ISG yielded 701,011 total SNP sites that were filtered to a subset of 304,497 SNP sites present in all of the genomes analyzed. These SNP sites were concatenated and used for phylogenetic analysis as previously described (50). A maximum-likelihood phylogeny with 1000 bootstrap replicates was generated using RAxML v.7.2.8 (51) and visualized using FigTree v.1.4.2 (http://tree.bio.ed.ac.uk/software/figtree/) and Interactive tree of life (52). Clades were assigned based on visual determination of groupings. Three genome outliers (1_176_05_S3_C2, 2_011_08_S1_C1, and 2_156_04_S3_C2 were removed from the tree figures for visualization purposes.

### Serotype identification

*In silico* serotype identification was performed on the assembled genomes using the online SerotypeFinder 1.1 (https://cge.cbs.dtu.dk/services/SerotypeFinder/) and an LS-BSR analysis using the serotype sequences compiled for the SRS2 program (https://github.com/katholt/srst2/tree/master/data) (15, 26).

### Multilocus sequence typing (MLST)

*In silico* MLST was performed on the assembled genomes using the Achtman *E. coli* MLST scheme (53). Gene sequences were identified in the isolate genomes using BLASTn and MLST profiles were determined by querying the PubMLST database (http://pubmlst.org).

### Variations in gene distributions

The gene content across all genomes was identified and compared using the large-scale BLAST score ratio (LS-BSR) with default settings, as previously described (27). Genes with a BSR value ≥0.80 are considered to be highly conserved and present in the isolate examined at this level of homology. Those genes that are conserved in all genomes were removed from further analyses. The predicted protein function of each gene cluster was determined using an ergatis-based (54) in-house annotation pipeline (55).

### Virulence factor and antibiotic resistance gene identification

The list of compiled common *E. coli* virulence factors genes was used for interrogation of the study genomes (Table S2). Antibiotic resistance genes were compiled from the Comprehensive Antibiotic Resistance Database (CARD; http://arpcard.mcmaster.ca, downloaded June 24, 2015) (28). The nucleotide sequence for each factor or resistance gene was aligned against all sequenced genomes with BLASTN (45) in conjunction with LS-BSR (27). Genes with a BSR value ≥0.80 were considered highly conserved and present in the isolate examined.

### Statistical analysis of macrolide resistance gene distributions

A logistic regression on the probability of a macrolide gene being present in an *E coli* isolate was run against 2 covariates: time point (excluding the baseline) or antibiotic treatment. For each individual, the two to three isolates were considered replicates for that time point, and the time points were far enough apart to be considered independent. Therefore, gene presence was collapsed as presence in at least one of the replicates at a given subject and time point. Each subject by time combination was considered an independent observation. Genes in this analysis with p-values ≤ 0.05 were considered significant. If the covariate was dichotomous, then the Wald Chi-Square test statistic was used to determine significance.

